## Supplementary material for "Multifaceted confidence in exploratory choice"

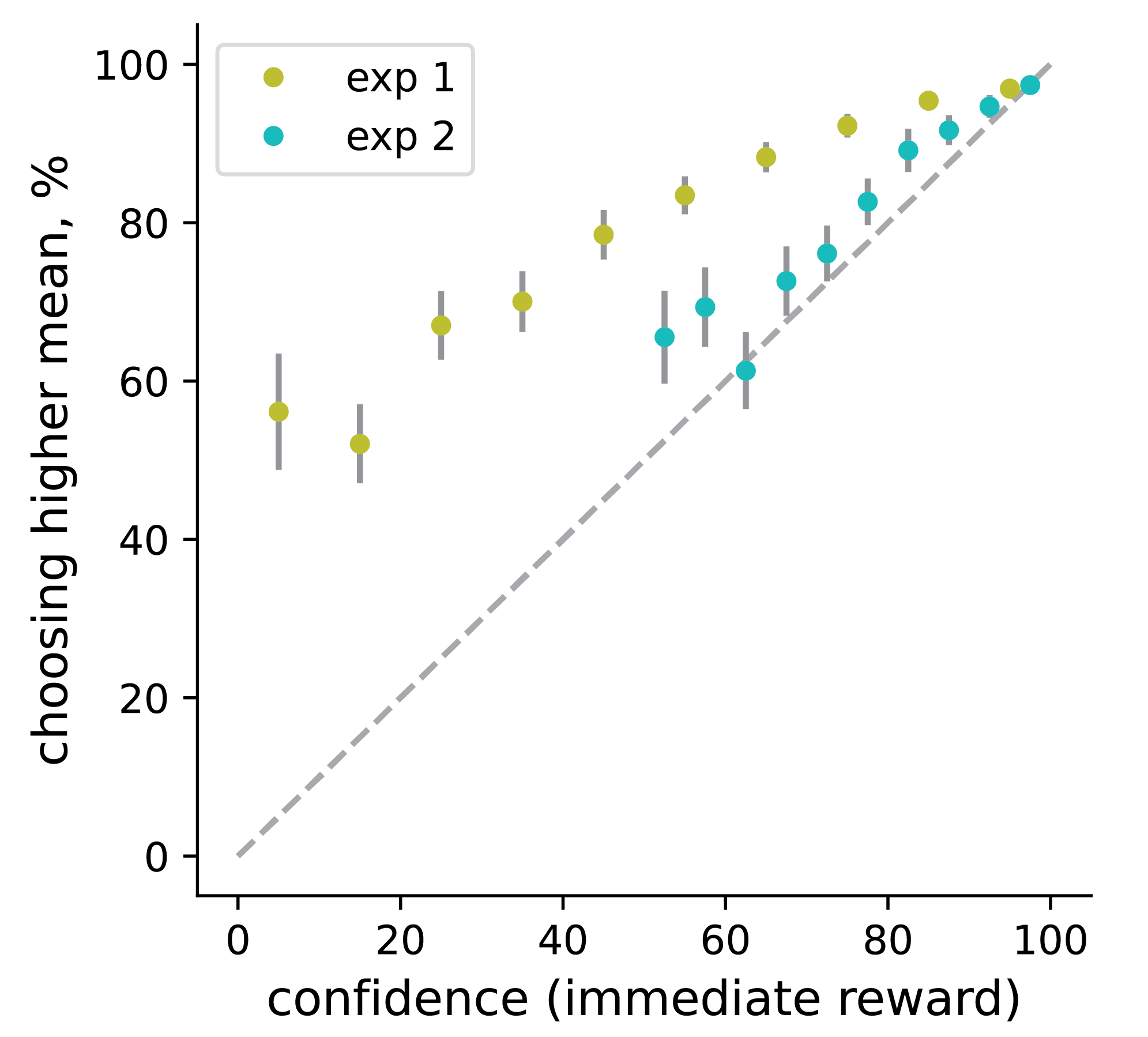

**Fig S1**. Confidence-accuracy calibration. Using half scale (50 to 100) to elicit confidence resulted in a better confidence-accuracy calibration. We binned the judgements about immediate reward, and for each bin, calculated the rate of choosing a higher mean option. The error bars represent the standard error of the mean. Judgements across both horizons were included, but focusing only on the short horizon produces a very similar result.

**S1_Appendix**

**Confidence questions**.

We piloted different versions of the questions about confidence. Initially, we used a 1-dimensional confidence scale, and pseudo-randomly presented on each trial one of 3 questions: 1) How confident are you that the choice you just made brings more points in this turn? 2) How confident are you that the choice you just made helps you collect more points in total in the current game? 3) How confident are you that the choice you just made was correct?

As we did not find a difference in confidence between these questions, we reasoned that subjects might have been confused about the exact question asked or, alternatively, just stop paying attention after some number of trials. We then switched to the 2-dimensional scale and asked two questions simultaneously: 1) How confident are you that the choice you just made leads to more points in turn 5? 2) How confident are you that the choice you just made leads to more points in total over turns 5-10? As this also did reveal a difference between confidence judgements, we adjusted the questions to the ones presented in the main text.

**Task instructions**.

We presented the participants with the following instructions, shown one or two sentences at a time and accompanied by relevant images of bandits or confidence scales:

"In this task, you will be playing a series of gambling games. In each game, you will be choosing between two slot machines like you might find in a casino. When you choose to play a particular machine, its payout will be shown. The machine you do not play on that turn will show XX. During a game, each slot machine will pay out a different number of points on average. The average of each machine will be randomly selected with equal probability to a number between 30 and 70. One of the machines will always pay more on average and will be the better option to choose in a game. However, it may be difficult to tell which is the better machine during a game. This is because, when played, a machine will give points with some (but the same for both machines) spread around its average. Importantly, both machines are sampled at each turn. You will get points in a turn only if the machine you chose had more points sampled than the one you didn't. This will not be signalled in each turn but will be reflected in the sum of points displayed after each game. Thus, the best strategy is to play the machine with a higher average. Your goal is to figure out the average worth of each machine, so you can choose the better machine and earn as many points as possible. During the experiment, there will be two types of games: 5-turn and 10-turn games. The number of turns is determined by the height of the machines. You will be told which machine to choose in the first 4 turns of each game. The green square will tell you which machine you MUST choose. For instance, here you would be asked to choose the left machine. After the first 4 turns, you will be free to choose between the two machines. The two green squares indicate that you are free to choose either machine. Following the first free choice, you will be asked to provide a confidence judgement about your choice. It is important that you pay attention to the question as one can be confident about different things. For example, if one was to make an investment, one would have to make a decision whether to consult a knowledgeable advisor for a small fee. One can be very confident that choosing to consult is the correct choice, while being somewhat less confident that his/her portfolio will double in the long term, and not being confident in a significant short term gain at all. You will be asked one or both of the following confidence judgements, on a scale from 0 [50 in Exp2] to 100 (0 = confident mistake, 50 = unsure, 100 = certain): 1. How confident are you that the choice you just made leads to more points in turn 5? 2. How confident are you that the choice you just made leads to more points in total over turns 5-10? In the example on the left, one is 100\% confident that their choice will lead to more points in turn 5, and 75\% confident that their choice will lead to more points in total over turns 5-10. Please click with your mouse and press 'Enter' to submit your response. Before submitting, you can adjust the response by dragging the pointer."

The participants were also instructed that their payment would depend on both choices and confidence judgement as follows:

"Your payment for taking part in this experiment is composed of 2 equal halves: 1. A computer program will choose a game you played randomly, and the points you earned will be converted to money. You can increase your chances to get a high payoff for this half if you collect a lot of points in as many games as possible. 2. A computer program will choose a game randomly, and reward you based on the confidence judgement you made in that game, using a lottery. The payoff of the lottery is set up in such a way that you can increase your payoff chances by evaluating and reporting your real confidence as accurately as possible in as many games as possible."
